# Online HD-tRNS over the right temporoparietal junction modulates social inference but not motor coordination

**DOI:** 10.1101/2025.04.14.648341

**Authors:** Quentin Moreau, Vincent Chamberland, Lisane Moses, Gabriela Milanova, Guillaume Dumas

## Abstract

Social interactions are fundamental to human cognition, with the right temporoparietal junction (rTPJ) playing a key role in integrating motor coordination and social inference. Transcranial random noise stimulation (tRNS) is a promising technique for modulating cortical excitability, yet its effect on dynamic social processes remains unexplored. This study aimed to establish a causal link between rTPJ and real-time social interaction by applying high-definition transcranial random noise stimulation (HD-tRNS) during the Human Dynamic Clamp (HDC) paradigm. Specifically, we investigated whether online HD-tRNS modulates motor coordination and social inference when interacting with an adaptive virtual partner (VP). Eighty right-handed participants were assigned to one of two experiments: (Exp1) a block design with active HD-tRNS and sham blocks or (Exp2) a trial-by-trial design with intermixed active and sham conditions. Participants engaged in an interactive coordination task with a covert VP that could behave cooperatively or competitively. Kinematic data and self-reported measures of perceived cooperativeness and humanness were analyzed. HD-tRNS over the rTPJ did not influence motor coordination or task performance. However, in Exp1, participants receiving active stimulation progressively attributed lower cooperativeness and humanness to the competitive VP, suggesting improved detection of competitive intent. This effect was absent in Exp2, indicating that repeated stimulation was necessary for cumulative neuromodulatory effects. Online HD-tRNS over the rTPJ does not acutely impact motor coordination but progressively modulates social inference, particularly in competitive interactions. The findings highlight the rTPJ’s role in self–other distinction and suggest that repeated short neuromodulation sessions can shape social perception.

**Significant statement:** Social interactions rely on distinguishing between cooperative and competitive behaviors, a process involving the right temporoparietal junction (rTPJ). Using high-definition transcranial random noise stimulation (HD-tRNS), we tested the rTPJ’s causal role in real-time social inference. While HD-tRNS did not affect motor coordination, repeated stimulation progressively reduced perceived cooperativeness and humanness of a competitive virtual partner, suggesting enhanced sensitivity to competitive intent. Our findings provide insights into social perception mechanisms and suggest HD-tRNS as a potential tool for studying and modulating social cognition, with implications for conditions affecting social inference, such as autism and schizophrenia.

## 1. Introduction

Social interactions are fundamental to human life, playing a key role helping individuals adapt to their sociocultural environment (Laland et al., 2001; Tomasello and Rakoczy, 2003). Understanding the neural correlates of such interactions is crucial for uncovering the mechanisms of social cognition. Early studies in social neuroscience relied on unidirectional paradigms in which isolated participants passively observed social stimuli, such as faces or other-directed actions (Hari and Kujala, 2009; Hari et al., 2015). At present, the use of Virtual Partners (VPs), which are computer-controlled avatars governed by computational models of human behavior (Kelso et al., 2009), enables experimenters to create ecological scenarios to study interpersonal interactions (Dumas et al., 2018). Combined with neuroimaging, they allow the examination of the neural underpinnings of real-time interaction in experimentally controlled environments (Fairhurst et al., 2013, 2014; Pfeiffer et al., 2014; Moreau et al., 2020, 2023; Formica and Brass, 2024).

Recently, the combination of high-density electroencephalography (EEG) with the Human Dynamic Clamp (HDC)—an interactive paradigm in which participants coordinate finger movements with a covert VP (Dumas et al., 2014b; Kelso et al., 2014)—highlighted the right temporoparietal junction (rTPJ) as a critical cortical region for integrating “low-level” motor coordination with “high-level” sociocognitive processes (Dumas et al., 2020). These findings align with extensive evidence recognizing the rTPJ as a key integrative hub in social cognition (Decety and Lamm, 2007; Lombardo et al., 2010; Bzdok et al., 2013; Carter and Huettel, 2013; Krall et al., 2015; Wu et al., 2015), mainly supporting functions related to the Theory of Mind (Saxe and Kanwisher, 2003; Schurz et al., 2017), self–other distinction (Decety and Sommerville, 2003; Sowden and Shah, 2014; Lamm et al., 2016; Quesque and Brass, 2019), and inhibition of motor imitation during social interaction (Brass et al., 2005, 2009; Spengler et al., 2009; Hogeveen et al., 2015; Sowden and Catmur, 2015).

However, while correlational studies provide strong evidence, they cannot alone establish the rTPJ’s causal role. A growing body of research has demonstrated that neuromodulating the rTPJ can influence various sociocognitive processes (Donaldson et al., 2015; Ahmad et al., 2021). Specifically, excitatory neuromodulation using high-frequency TMS or anodal transcranial direct current stimulation (tDCS) has been shown to enhance self–other distinction (Santiesteban et al., 2012a, 2015), perspective-taking (Martin et al., 2020), and intention attribution (Schuwerk et al., 2021; Panico et al., 2024). Conversely, inhibitory protocols such as low-frequency TMS or cathodal tDCS reliably disrupt these processes (Mai et al., 2016; Bardi et al., 2017; Coll et al., 2017; Era et al., 2020). However, the predominant use of offline protocols has constrained our understanding of how online neuromodulation might shape ongoing, dynamic social interaction.

Transcranial random noise stimulation (tRNS) represents a relatively new tES technique that delivers random current fluctuations within a broad frequency band to modulate cortical excitability (Terney et al., 2008). Although neurophysiological mechanisms of tRNS are still under investigation, it is hypothesized to enhance cortical excitability via stochastic resonance, thereby improving the signal-to-noise ratio in targeted neural areas (McDonnell and Abbott, 2009; Van Der Groen and Wenderoth, 2016; Battaglini et al., 2023). Importantly, short sessions of tRNS have been shown to induce acute changes in neuronal activity and behavior (Potok et al., 2021, 2022), making this technique particularly promising for online neuromodulation during interactive paradigms (Cappelletti et al., 2013; Pirulli et al., 2013; Van Der Groen et al., 2022). In this study, we hypothesized that tRNS over the rTPJ would enhance participants’ motor coordination with the VP and modulate their social inference of the VP’s cooperativeness and humanness, consistent with the proposed integrative role of the rTPJ in bridging low- and high-level cognitive processes. Two complementary experiments were carried out: Experiment 1 investigated the acute effects of online tRNS on participants’ coordination and social inference across randomized blocks, and Experiment 2 systematically probed the consistency and specificity of possible acute effects across randomized trials.

## 2. Materials and methods

### 2.1. Participants

Eighty French-speaking, right-handed participants with normal or corrected-to-normal vision, no prior history of neuropsychiatric conditions or movement disorders, and no contraindication to tES (Bikson et al., 2016; Antal et al., 2017) were recruited for the study. Forty participated in Experiment 1 (24 women; mean age = 29.6, SD = 11.7) and forty in Experiment 2 (25 women; mean age = 32.6, SD = 13.5). One participant from Experiment 1 was excluded due to non-compliance with instructions. Participants provided written informed consent and received $30 CAD compensation. The experiments lasted approximately three hours (Experiment 1) or two hours (Experiment 2). The study protocol was approved by the Ethics Review Board of the CHU Sainte-Justine (September 19, 2022) and complied with the Declaration of Helsinki.

### 2.2. Procedure

Upon arrival, participants were seated, fitted with the tRNS headset, and introduced to the HDC paradigm. They were informed that some trials would involve interacting with a human partner in another room, while others could feature a virtual partner designed to imitate human-like behaviors. To minimize inter-individual variability in the quality of script execution, the same researcher conducted all interventions. Participants in Experiment 1 completed two blocks of 40 trials separated by a one-hour washout period to minimize a potential carry-over effect of the stimulation, while participants in Experiment 2 completed a single block comprising a pseudo-randomized mix of 20 active and 20 sham trials (Fig. 1B). Potential tRNS adverse effects were assessed at the end of each block (see Supplementary material). At the end of the procedure, participants were informed that no human was involved and completed a Turing-like test, rating their belief (0–100%) in having interacted with a human.

**Figure 1.**
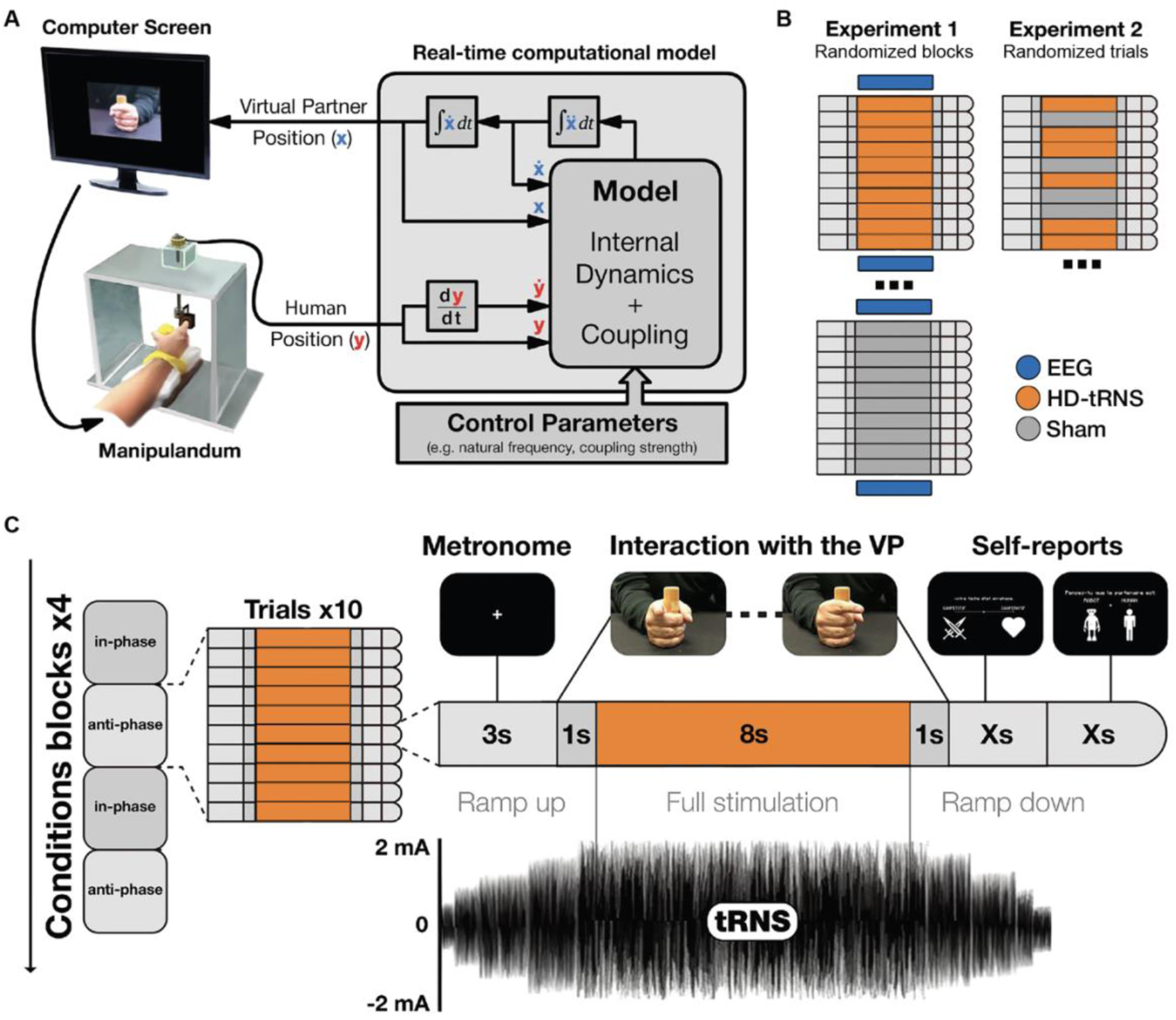
(A) The Human Dynamic Clamp (HDC) interactive paradigm uses a real-time computational model to dynamically generate the VP’s movement. (B) The two experimental designs: Experiment 1 with EEG recordings and randomized active stimulation and sham blocks; and Experiment 2 with randomized active stimulation and sham trials. (C) Example of a typical HDC trial with active stimulation.

### 2.3. Task

The HDC paradigm (Dumas et al., 2014a) consists of an experimental design that captures participants’ finger kinematics in real-time, which then feeds continuously into a nonlinear dynamical model (the Haken-Kelso-Bunz model of Coordination Dynamics [Haken et al., 1985]) driving the VP’s finger movements on a monitor (Fig. 1A). The goal of the task is to be as synchronous as possible with the finger movements of the VP, either in an imitative (in-phase) or complementary (anti-phase) fashion. Each trial started by indicating the intended coordination pattern (in-phase or anti-phase). Participants then entrained their movements to a 1.6 Hz auditory metronome before the VP’s finger appeared, after which they maintained the same frequency and attempted the specified coordination pattern (Fig. 1C). The VP was configured to align cooperatively or competitively, either adaptively supporting or opposing the participant’s intended coordination pattern. Each trial yielded coordination measures: Motor (movement similarity), Coordination (phase alignment), and Task (goal alignment: in-phase or anti-phase). Immediately following the interaction phase, participants self-reported whether they perceived the partner as cooperative or competitive on a continuous scale, and if they believed they were interacting with a human or a computer in a binary choice. See Supplementary material for detailed HDC setup and scores.

### 2.4. HD-tRNS montage

A Starstim8® wireless EEG-tES device (Neuroelectrics, Barcelona, Spain) was used to apply both tRNS and EEG recordings (EEG montage, procedure, analysis, and results are presented in Supplementary material). Five Ag/AgCl hybrid electrodes (1 cm radius, π cm² contact area) were arranged in a ring-like (4 × 1) high-definition tRNS (HD-tRNS) montage on a neoprene head cap following the international 10-10 system. The central electrode targeting the rTPJ was placed at CP6 based on consensus in previous tDCS studies (Santiesteban et al., 2015, 2012b; Sowden et al., 2015; Vandenbroucke et al., 2016; Esse Wilson et al., 2018; Martin et al., 2020; Wu et al., 2023; Cesari et al., 2024) and delivered or received 100% of the current, while four peripheral electrodes (C4, T8, P4, and P8) each received or delivered 25% (Fig. 2B). The applied current was approximately 2 mA (1935 µA, peak-to-baseline) with no offset, sampled at 1280 Hz with a standard deviation of 645 µA, and drawn from the full spectrum of the available high-frequency range of 100–500 Hz to maximize cortical excitability (Moret et al., 2019).

**Figure 2.**
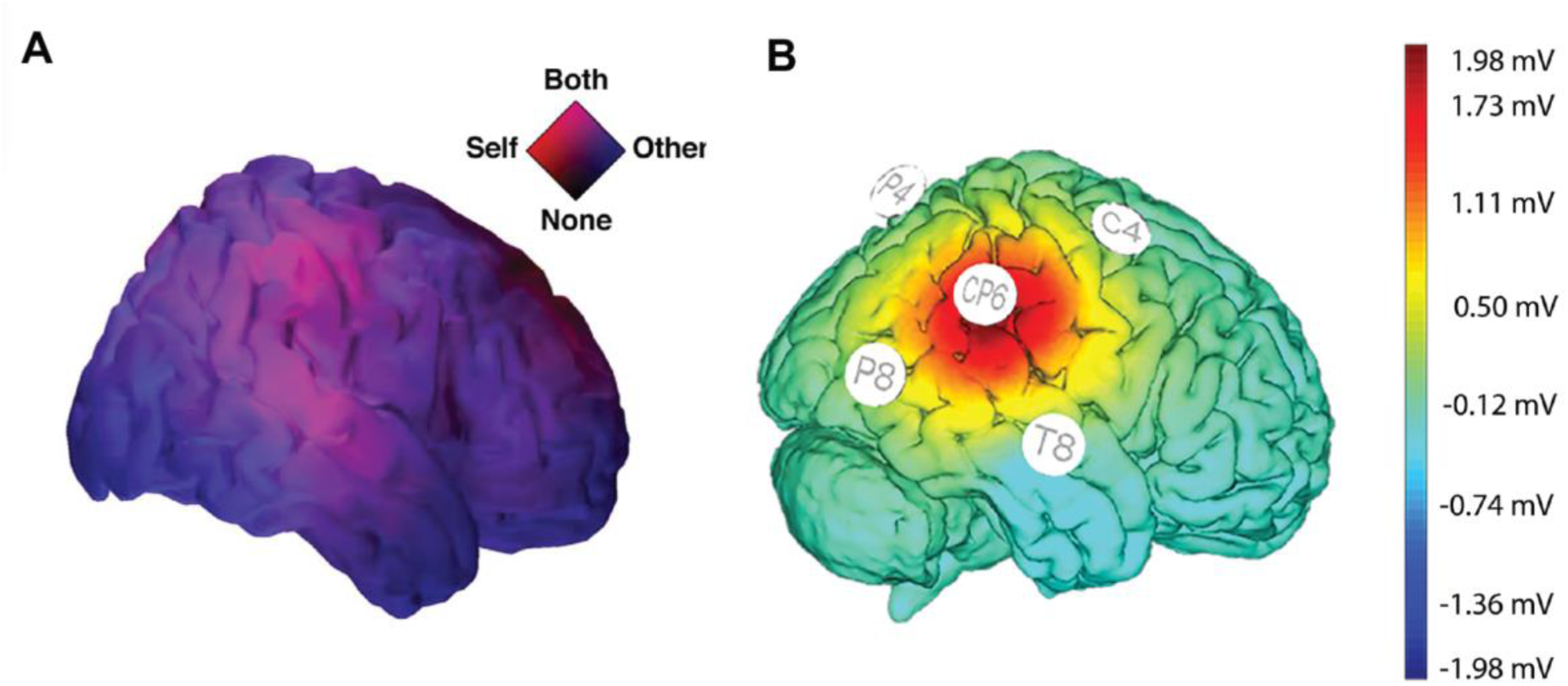
(A) Estimated cortical sources of high-density EEG revealing the rTPJ is involved in the integration of self and other during the HDC paradigm, adapted from Dumas and al. (2020). (B) Simulated focal stimulation of the rTPJ using HD-tRNS modeled in NIC 2.0 for the current experiment.

HD-tRNS was administered during interaction with the VP to deliver acute effects (Fig. 1C). The HDC original code was modified and combined with a custom MatLab® script to interact with the Neuroelectrics Instrument Controller (NIC 2.0) software to load and start the active or sham stimulation protocol at the beginning of each HDC trial. During each active trial, the current would ramp up for four seconds, be maintained for eight seconds during the interaction with the VP, and then ramp down for four seconds, for a total of 16 seconds. In the sham trials, the current would only ramp up and down. This cycle delivered about 7.7 mC to the scalp per trial in the active condition, whereas the sham condition delivered about 4.8 mC. Participants received a cumulative dose of approximately 500 mC in Experiment 1 or 250 mC in Experiment 2.

**Supplementary Figure 1.**
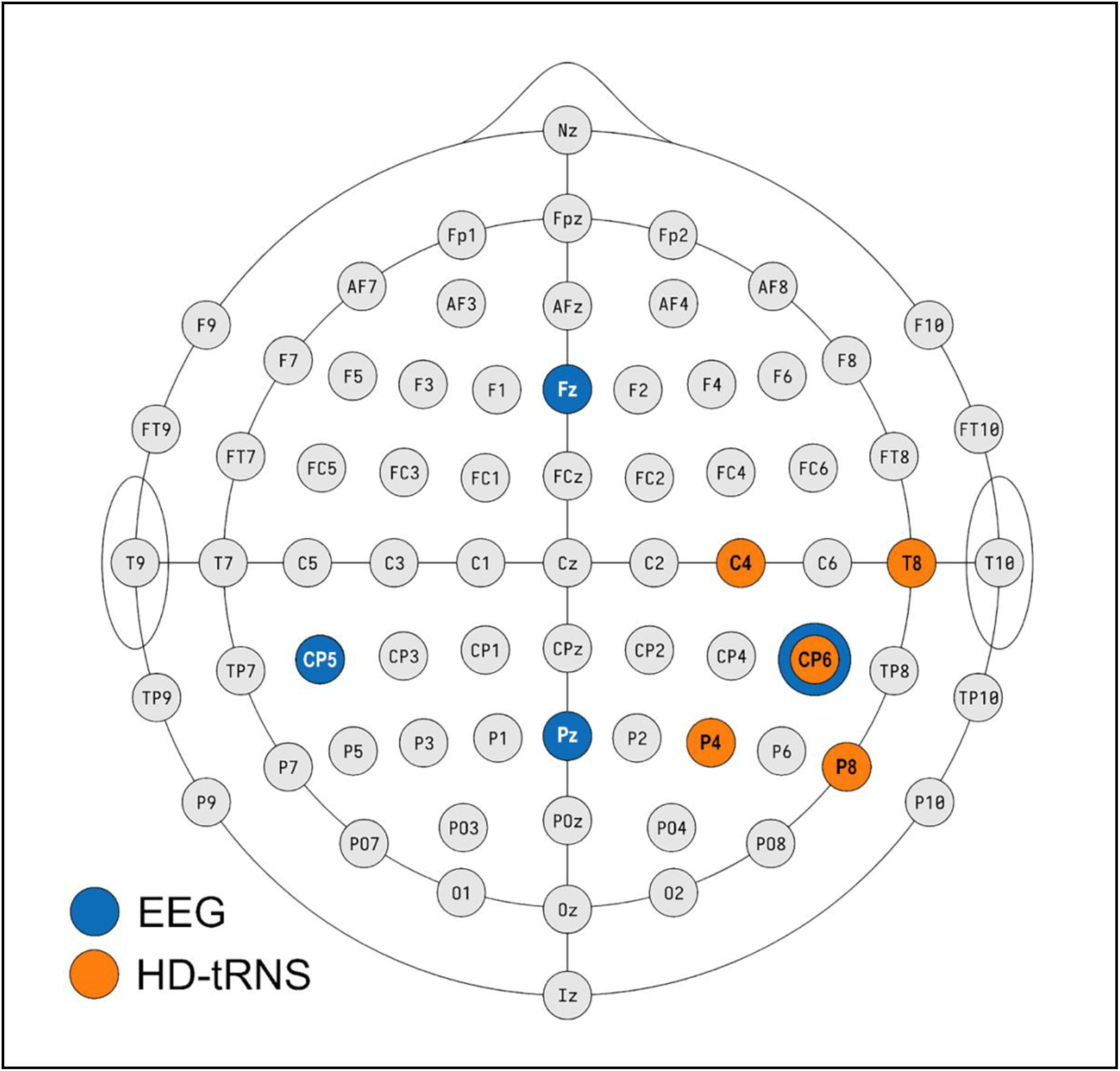
Electrode placement for the hybrid HD-tRNS and EEG setup.

### 2.6. Data and statistical analysis

Using Python (v3.12.3), we imported the raw kinematic data (participant and VP positions and velocities) and corrected outliers in the participant’s movement trajectories via a DBSCAN clustering algorithm (scikit-learn v1.6.0 [Pedregosa et al., 2011]). We then mean-centered the cleaned data and applied a second-order, double-pass Butterworth filter with a 20 Hz cutoff (scipy v1.14.1 [Virtanen et al., 2020]) to reduce high-frequency noise. Finally, we computed the continuous relative phase between the participant’s and the VP’s movements using a Hilbert transform to estimate phase and amplitude required for motor coordination scores. Attribution of Cooperativeness and Humanness scores were directly retrieved from the trial reports using participants’ self-assessments.

All statistical analyses were performed in R (v4.3.3) using generalized linear mixed models (GLMMs) with *glmmTMB* (v1.1.10 [Brooks et al., 2017]). For Coordination, Motor, Task, and Cooperativeness attribution scores (bounded 0–1), we applied Beta regression with a logit link; for binary Humanness attribution scores, we used a binomial GLMM with a logit link. Stimulation protocol (active/sham) and VP behavior (cooperative/competitive) were included as fixed effects, with random intercepts for participants to capture individual baseline differences. Trial number (z-scored) was added as a continuous fixed effect to account for cumulative stimulation or learning. Fixed effects were tested via Wald z-tests, and estimated marginal trends were obtained and compared using *emmeans* (v1.10.5 [Lenth, 2025]) with Tukey-adjusted pairwise comparisons. Exponentiated coefficients (odds ratios) and 95% confidence intervals are reported. The results of the questionnaires on adverse effects were analyzed using a MANOVA on all criteria (itching, pain, burning, heat, metal taste and fatigue).

## 3. Results

In line with previous HDC studies, participants mainly believed that another human took part in the experiment as an interacting partner: 72.22% (SD = 24.76) in Experiment 1 and 69.05% (SD = 28.67) in Experiment 2The MANOVA results indicated no significant difference in adverse effects between the active and sham stimulation conditions (Wilks’ λ = 0.9501, F(5,74) = 0.78, p = 0.57). This suggests that, based on these implicit indicators, participants were unable to distinguish between active and sham stimulation blocks.

### 3.1. Online HD-tRNS does not modulate motor coordination and task performance

In Experiment 1, no main effect of HD-tRNS was found on Coordination (Exp(β) = 0.96 [0.90, 1.02], p = 0.221), Motor (Exp(β) = 1.00 [0.95, 1.04], p = 0.880) and Task scores (Exp(β) = 1.02 [0.97, 1.08], p = 0.483), and no significant interactions were observed. Similarly, in Experiment 2, active stimulation yielded no significant main effect on Coordination (Exp(β) = 0.93 [0.85, 1.02], p = 0.141), Motor (Exp(β) = 0.99 [0.93, 1.06], p = 0.833) and Task scores (Exp(β) = 1.01 [0.95, 1.08], p = 0.785), and no significant interactions emerged. These results suggest that online HD-tRNS over the rTPJ could not modulate any level of motor coordination in any experiment and did not affect task performance.

### 3.2. Online HD-tRNS progressively improves recognition of competitive behavior

In Experiment 1, despite no main effect of HD-tRNS on cooperativeness attribution scores (Exp(β) = 1.00 [0.88, 1.13], p = 0.996), a significant three-way interaction was found between stimulation protocol, VP behavior, and trial progression (Exp(β) = 1.56 [1.31, 1.84], p < 0.001). Post-hoc trend analyses further showed that, only when faced with a competitive VP, active stimulation led to a significant decrease of perceived cooperativeness (Exp(β) = 0.72 [0.66, 0.80], p < 0.001), with the contrast between sham and active stimulation being also significant (Exp(β) = 1.47 [0.25, 0.52], p < 0.001). Conversely, when the VP was cooperative, both active stimulation (Exp(β) = 1.16 [1.08, 1.24], p < 0.001) and sham (Exp(β) = 1.09 [1.02, 1.17], p = 0.017) were associated with increasing cooperativeness attribution over time, but the difference between these two protocols was not significant (Exp(β) = 0.94 [0.85, 1.04], p = 0.231).

In Experiment 2, despite another non-significant main effect of HD-tRNS (Exp(β) = 1.02 [0.86, 1.21], p = 0.780), a three-way interaction also emerged between stimulation protocol, VP behavior, and trial progression (Exp(β) = 0.77 [0.61, 0.98], p = 0.034). However, contrary to results from Experiment 1, post-hoc trend analyses revealed that active stimulation had no significant effect on cooperativeness attribution in the competitive VP condition (Exp(β) = 1.12 [0.98, 1.28], p = 0.084), and the contrast between both protocols was not significant (Exp(β) = 0.95 [0.79, 1.14], p = 0.059). When the VP was cooperative, only sham had a significant upward trend in cooperativeness attribution over time (Exp(β) = 1.28 [1.42, 1.44], p < 0.001). The contrast between both protocols was also significant (Exp(β) = 1.23 [1.05, 1.44], p = 0.009).

These findings indicate that only in the randomized block design of Experiment 1 did active stimulation alter cooperativeness attribution over trials during competitive interactions with the VP (Fig. 3), suggesting that online HD-tRNS over the rTPJ progressively improves participants’ ability to detect competitive cues and accurately interpret the VP’s behavior.

**Figure 3.**
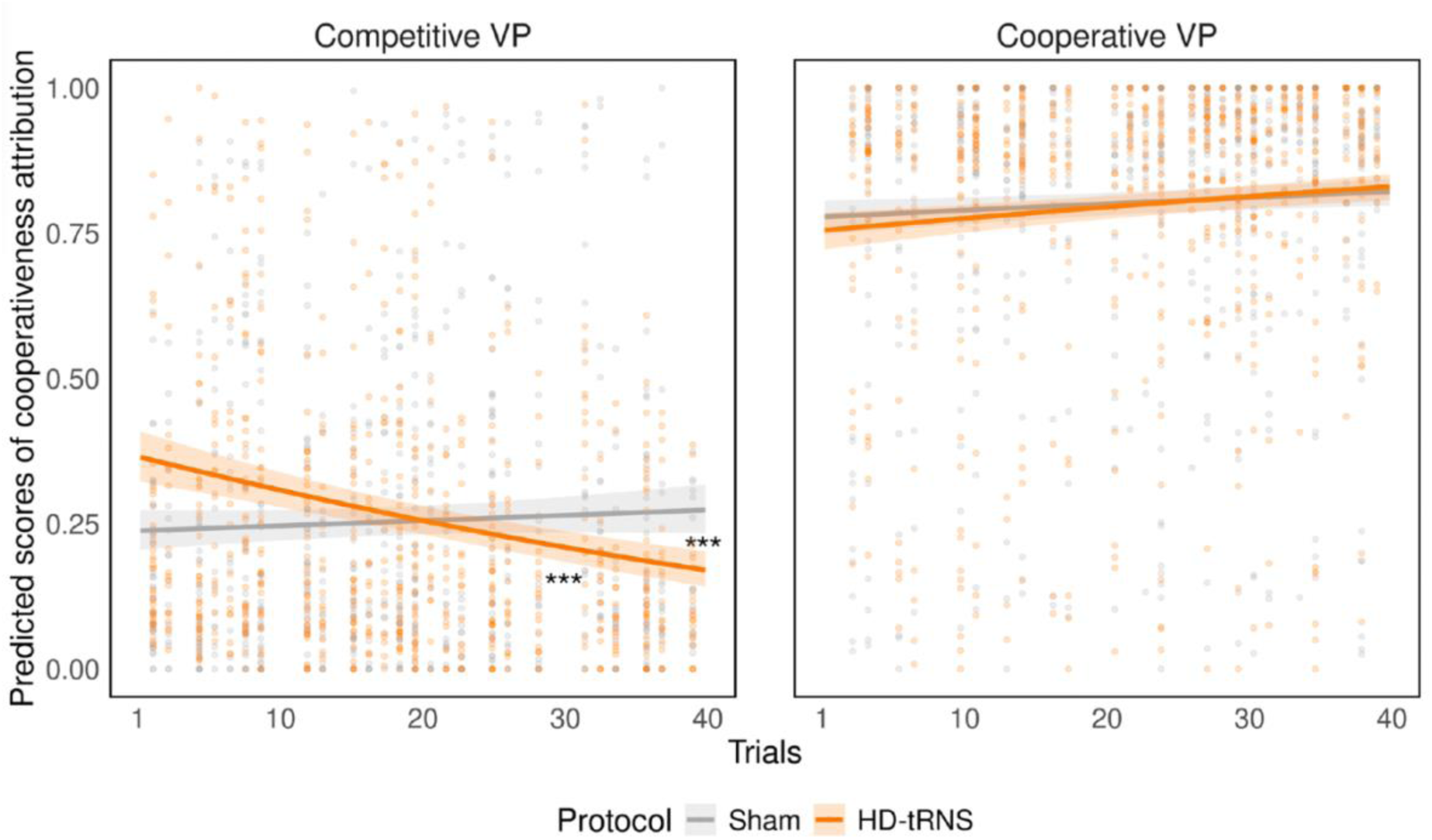
Predicted scores of cooperativeness attribution in Experiment 1 across stimulation protocols, VP behaviors, and trials. Scores are converted back from the logit scale to the original probability scale for visualization.

### 3.3. Online HD-tRNS progressively reduces humanness attribution in competitive interactions

In Experiment 1, HD-tRNS showed a significant main effect, indicating lower humanness attribution scores under active stimulation compared to sham at baseline (Exp(β) = 0.72 [0.58, 0.90], p = 0.004), and a significant three-way interaction was found between stimulation protocol, VP behavior, and trial progression (Exp(β) = 1.48 [1.09, 2.02], p = 0.012). Post-hoc trend analyses showed that when interacting with a competitive VP, only active stimulation resulted in a significant decrease in humanness attribution (Exp(β) = 0.74 [0.62, 0.89], p = 0.001), with the contrast between sham and active stimulation also being significant (Exp(β) = 1.50 [1.17, 1.93], p = 0.001). In comparison, when the VP was cooperative, both active stimulation (Exp(β) = 0.86 [0.76, 0.98], p = 0.025) and sham (Exp(β) = 0.87 [0.77, 1.00], p = 0.042) showed significant decreasing humanness attribution over time, and the contrast between protocols was not significant (Exp(β) = 1.01 [0.84, 1.22], p = 0.890).

Furthermore, exploratory analysis revealed a significant interaction between stimulation protocol and baseline humanness attribution extracted from the random intercepts (Exp(β) = 1.78 [1.22, 2.60], p = 0.003), indicating that the effect of HD-tRNS weakens with increasing baseline humanness attribution. Post-hoc analyses showed that at lower baseline levels (logit = -0.681), HD-tRNS significantly reduced humanness attribution compared to sham (Exp(β) = 0.54 [0.41, 0.76], p < 0.001). However, at higher baseline levels (logit = 0.448), there was no significant difference between active stimulation and sham (Exp(β) = 0.89 [0.75, 1.06], p = 0.347). These findings demonstrate that HD-tRNS had a greater effect on reducing humanness attribution for participants with lower baseline humanness attribution, while its effect diminished as baseline levels increased.

In Experiment 2, HD-tRNS did not produce a significant main effect (Exp(β) = 1.02 [0.77, 1.36], p = 0.884), and no significant interactions were observed. Together, these findings demonstrate that only within the randomized block design of Experiment 1 did active stimulation modulate changes in humanness attribution over trials during competitive interaction with the VP (Fig. 4A), suggesting that online HD-tRNS over the rTPJ progressively reduces the perceived humanness of a competitive partner, particularly in individuals with lower baseline humanness attribution (Fig. 4B).

**Figure 4.**
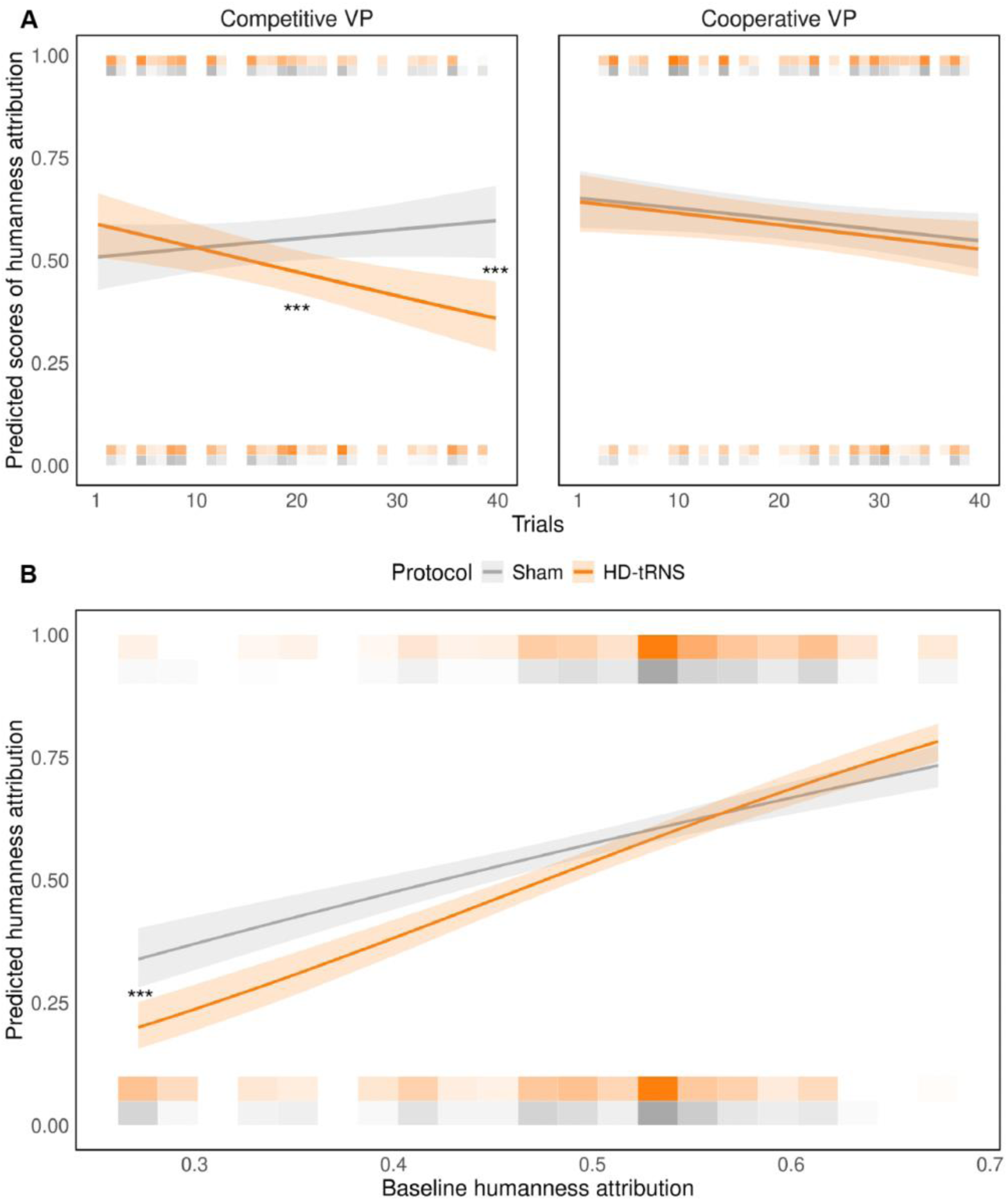
Predicted scores of humanness attribution by stimulation protocols in Experiment 1, (A) cross VP behaviors and trials, and (B) compared to baseline humanness attribution. Scores are converted back from the logit scale to the original probability scale for visualization.

## 4. Discussion

Our study sought to investigate the causal contribution of the rTPJ to real-time social interaction via HD-tRNS during the HDC paradigm. We found that while short, repeated sessions of online HD-tRNS did not acutely or cumulatively alter motor coordination with the VP, they progressively modulated participants’ attributions of competitiveness and humanness, revealing a potential acute role of the rTPJ excitability in social inference.

### 4.1. No modulation of low-level motor coordination

Previous research had linked stimulation of rTPJ with modulatory effects on motor coordination. Notably, Era and al. (Era et al., 2020) demonstrated that inhibiting the rTPJ with offline continuous theta burst stimulation selectively disrupted imitative interactions with a VP. Our results show no modulatory effect of HD-tRNS over the rTPJ on any motor coordination scores. One possible explanation is that while rTPJ can indirectly influence motor imitation-inhibition (Brass et al., 2005, 2009; Hogeveen et al., 2015; Sowden and Catmur, 2015), its role in simple in-phase or anti-phase coordination may be less critical. Additionally, it is possible that our participants were already performing at a ceiling level regarding motor skills, leaving little room for tRNS to exert a measurable effect on these variables. Finally, the discrepancy between previous findings and ours may stem from differences in stimulation protocols. Future studies should further explore the conditions under which rTPJ stimulation influences motor coordination, particularly in tasks requiring greater self–other integration.

### 4.2. Modulation of high-level social inference requires cumulative stimulation

Experiment 1 demonstrated that HD-tRNS over the rTPJ modulated participants’ social inference. Only when interacting with a competitive VP across an active stimulation block did participants progressively attribute less cooperativeness and lower humanness to their partner. Interestingly, no effects were observed in Experiment 2, which intermixed active and sham protocols at the single-trial level.

Specifically, under active stimulation, participants became progressively more accurate in recognizing the VP’s covertly competitive behaviors, suggesting that excitatory rTPJ stimulation selectively strengthens self–other distinction in adversarial scenarios. This finding is consistent with neuromodulation studies that have either enhanced the rTPJ activity to improve its engagement in self–other distinction and perspective-taking tasks (Santiesteban et al., 2012b, 2015; Zhang et al., 2019; Martin et al., 2020) or inhibited the rTPJ activity, thereby impairing these functions (Tsakiris et al., 2008; Wang et al., 2016; Era et al., 2020). One prominent view is that the rTPJ dynamically directs attention to salient social cues, especially in ambiguous or adversarial interactions, evidenced by heightened rTPJ activity during competition compared to cooperation (Bitsch et al., 2018). Because competition often requires anticipating and countering an opponent’s goals, robust self–other distinction becomes critical for accurately separating one’s actions from an adversary’s (Giardina et al., 2011), and enhanced rTPJ excitability may tip individuals toward more precise recognition of competitive intent.

We also observed that active stimulation progressively lowered participants’ attribution of humanness toward the covertly competitive VP. By amplifying self–other distinction, the rTPJ may improve the identification of an adversary’s antagonistic intentions and thereby down-regulate empathic engagement or anthropomorphism (Payne and Tsakiris, 2017; Filmer et al., 2019). Notably, while the rTPJ is implicated in recognizing others as intentional agents (Özdem et al., 2017; Ogawa and Kameda, 2019), inhibiting its function has been shown to impair mentalizing and cognitive empathy (Mai et al., 2016; Coll et al., 2017). These results also align with evidence that perceiving interactions as more cooperative or synchronized increases empathy and attribution of humanness towards a covert partner (Koehne et al., 2016; Jastrzab et al., 2024), whereas adversarial or unstable interactions can diminish these impressions (Zhang et al., 2016; Dumas et al., 2020). During the HDC paradigm, the competitive behavior of the VP may have amplified this dissociative effect, with rTPJ stimulation enhancing self–other distinction and prompting participants to perceive the antagonistic VP as less human.

Importantly, our exploratory analysis revealed that the impact of HD-tRNS on humanness attribution was modulated by participants’ baseline perceptions of the VP. Specifically, the stimulation effect was most pronounced for participants with lower baseline humanness attribution, as HD-tRNS significantly reduced ratings in this group compared to sham. In contrast, the effect diminished when participants initially rated the VP more human-like. This interaction suggests that rTPJ stimulation may have a ceiling effect on humanness attribution, where participants who already perceive the partner as highly human show less room for further reductions. These findings align with models positing that the rTPJ integrates external cues (such as a partner’s uncooperative behavior) with internally generated predictions (Lamm et al., 2007; Carter and Huettel, 2013). For participants with lower baseline humanness attribution, stimulation may enhance sensitivity to antagonistic cues, reinforcing their perception of the partner as less human. Conversely, participants with higher baseline humanness attribution might already have robust social appraisals that resist significant shifts under stimulation. This baseline-dependent modulation highlights the rTPJ’s role in dynamically calibrating social perceptions and underscores the importance of individual differences in shaping neuromodulation outcomes.

A key takeaway is that only the randomized block design in Experiment 1 produced significant, cumulative rTPJ effects, whereas the single-trial randomization in Experiment 2 did not. This contrast suggests that repeated short HD-tRNS sessions in Experiment 1 allowed neuromodulatory effects to build up, whereas frequent switching in Experiment 2 likely disrupted progressive rTPJ strengthening. Prior research indicates that at least four to seven minutes of offline tRNS can yield robust behavioral outcomes (Chaieb et al., 2009, 2011; Haeckert et al., 2020), and one study applying tRNS to the rTPJ found that effects on temporal attention peaked around 15 minutes of stimulation (Tyler et al., 2018). Although brief, seconds-long tRNS sessions have been shown to induce rapid excitability changes in lower-level perceptual or motor tasks (Van Der Groen and Wenderoth, 2016; Van Der Groen et al., 2019; Potok et al., 2021, 2022), higher-level sociocognitive processes may require steadier or repeated stimulation to surpass the threshold for measurable effects. Overall, these findings underscore that repeated, trial-based HD-tRNS can progressively modulate complex social cognition, provided it is delivered in a context that allows incremental buildup without frequent protocol switching.

### 4.3. Limitations

Our study has several limitations. First, although we used a high-definition tRNS montage to target the rTPJ, the region’s histological heterogeneity (encompassing the angular and supramarginal gyri [Bzdok et al., 2013; Krall et al., 2015; Doricchi, 2022]) may have led to partial stimulation of neighboring subregions, reducing specificity. Second, while participants were unaware of their stimulation protocol, the experiment was not double-blinded, as experimenters knew the active or sham conditions, possibly introducing subtle bias despite standardized procedures. Third, we did not include a control stimulation site (e.g., the left TPJ, which has also been implicated in social inference and mentalizing [Ogawa and Kameda, 2019; Hao et al., 2022; Golec-Staśkiewicz et al., 2022]), limiting our ability to attribute effects solely to the rTPJ. Finally, we did not assess individual responses to tRNS before the experiment, potentially introducing inter- and intra-individual variability (López-Alonso et al., 2014) that may have influenced stimulation efficacy and contributed to variability in our results.

## 5. Conclusion

Our findings demonstrate that short online HD-tRNS sessions over the rTPJ do not affect basic motor coordination with a VP but significantly influence the dynamic process of social inference, particularly in competitive contexts. These effects appeared cumulatively over repeated trials within the randomized block design, suggesting a build-up of neural changes under sustained rTPJ stimulation. Together, the progressive shifts in how participants judged the VP’s cooperativeness and humanness underscore the rTPJ’s causal role in integrating self– other representations during competitive interactions.

Future research could explore closed-loop stimulation protocols to assess dynamically the causality of neuromodulatory effects (Ramot and Martin, 2022), as well as broader network-level modulation, potentially involving multi-site stimulation of interconnected regions such as the rTPJ and right-lateralized fronto-parietal areas to engage the distributed neural mechanisms underpinning social interaction (Dumas et al., 2020). Multi-brain stimulation targeting the rTPJs of interacting participants is another promising technique analogous to hyperscanning to study and manipulate inter-brain synchronization during social interaction (Dumas, 2022). Investigating these approaches in clinical and developmental populations with atypical social cognition, such as individuals with autism spectrum disorder and schizophrenia, may inform the design of adaptive precision neuromodulation interventions (Medaglia et al., 2020). Collectively, these directions have the potential to deepen our understanding of the rTPJ’s involvement in social interaction and expand the therapeutic applications of targeted neuromodulation.

## Acknowledgements

The authors would like to thank Dr. Hugo Théoret for lending the neuromodulation device, Dr. Karim Jerbi for his theoretical insights, and the Neuroelectrics team for their technical assistance.

## CRediT authorship contribution statement

**Quentin Moreau:** Conceptualization, Methodology, Formal analysis, Investigation, Writing – Original draft, Writing – Review & Editing, Visualization, Supervision, Project administration, Funding acquisition; **Vincent Chamberland:** Methodology, Formal analysis, Investigation, Writing – Original draft, Writing – Review & Editing, Visualization, Data curation, Project administration, Funding acquisition; **Lisane Moses:** Investigation, Data curation, Project administration; **Gabriela Milanova:** Investigation, Data curation, Project administration; **Guillaume Dumas:** Conceptualization, Methodology, Formal analysis, Writing – Review & Editing, Visualization, Supervision, Resources, Funding acquisition.

## Data availability

Data preprocessing and statistical analysis scripts, the modified HDC code, and the custom MatLab® script used to load stimulation protocols are available on our laboratory’s GitHub repository (https://github.com/ppsp-team/CLONE).

## SUPPLEMENTARY MATERIAL

### 1. HDC information

#### 1.1. Setup

Each participant was seated at a table in front of a 27-inch monitor (2560 × 1440 pixels, 144 Hz) positioned ∼60 cm away, with the right forearm placed on a U-shaped support (21.5 × 8 × 4 cm) and the right hand grasping a vertical Plexiglas cylinder (4.5 × 3 cm), leaving only the index finger free to move. The hand was oriented in the sagittal plane, and the distal part of the index finger was inserted into the circular orifice (2 cm diameter) of a Plexiglas block connected to a freely rotating metallic stem (18 cm in length). The angular displacement of the stem was recorded by a linear potentiometer, and this setup—fixed at the top of a Plexiglas box (30.5 × 31.5 × 20 cm) positioned ∼50 cm to the right of the participant’s midline—allowed friction-free, horizontal-plane flexion-extension about the metacarpophalangeal joint.

#### 1.2. Scores

**Coordination score**: quantifies the phase alignment between the human participant and the virtual partner (VP). It is calculated as the mean of the complex exponential of the phase difference (Δϕ) between the human and the VP, providing a synchronization measure. This score ranges from 0 to 1, where higher values indicate stronger phase coupling and better temporal coordination.

**Motor score**: measures the morphological similarity between the human and VP movements by comparing their amplitudes. It is derived using a normalization formula that assesses the relative difference and sum of their amplitudes over the trial duration. Values closer to 1 indicate high morphological similarity, while values near -1 suggest a divergence in movement dynamics.

**Task score**: evaluates how closely the participant’s performance aligns with the task goal, such as maintaining in-phase (0) or anti-phase (*π*) coordination with the VP. It is computed by assessing the normalized deviation of the phase difference from the target phase relationship. A Task score of 1 indicates perfect adherence to the task goal, while lower scores reflect increasing deviation.

**Cooperativeness score**: evaluates the participant’s ability to accurately attribute intention to the VP’s behavior as either “cooperative” or “competitive.” Derived from the trial self-report, this score ranges from 0 (fully competitive) to 100 (fully cooperative) and provides a measure of how effectively the participant perceives and interprets the VP’s interaction style. It offers a window into the participant’s cognitive processing of social cues and their ability to discern the VP’s intended behavior.

**Humanness score**: represents the participant’s binary classification of the VP as either robotic (0) or human (1). This score is directly extracted from the trial self-report and reflects the participant’s categorical perception of the VP’s behavior based on its naturalness and anthropomorphic qualities.

### 2. EEG

#### 2.1. Procedure

In Experiment 1, each block (active or sham stimulation) was preceded and followed by 3 minutes of eyes-opened resting EEG recording (Fig. 1B). In Experiment 2, no pre- and post-block EEG was recorded.

#### 2.2. EEG Montage

The EEG montage consisted of three hybrid electrodes not included in the HD-tRNS montage (CP5, Fz, and Pz), along with one electrode also used for stimulation (CP6), forming a low-definition yet symmetrical setup (Supplementary Fig. 1). The reference electrode was positioned on the right earlobe, and the recording was conducted at a sampling frequency of 500 Hz.

#### 2.3. Data and statistical analysis

Each electrode (CP5, CP6, Fz, and Pz) of the eyes-open resting EEG recorded before and after every block of Experiment 1 was processed in the *MNE-Python* environment (v1.9.0; Gramfort et al., 2013). Data were band-pass filtered from 1–100 Hz, subjected to a 60 Hz notch filter, and non-neural artifacts were automatically identified and removed using the included *iclabel* algorithm (Pion-Tonachini et al., 2019). The cleaned signals were segmented into 2-second epochs, and residual artifacts were addressed with the *autoreject* algorithm (v0.4.3; Jas et al., 2017). Power spectral densities were calculated for each condition using Welch’s method with 1 Hz resolution, and aperiodic (1/f-like) and periodic (oscillatory) components of the spectra were parameterized using the *specparam* algorithm (legacy *fooof* v1.1; Donoghue et al., 2020).

To analyze the data, linear mixed models (LMMs) via *statsmodels* (v0.14.4; Seabold and Perktold, 2010) were used to assess how the differences between the four experimental conditions (pre/post-active, pre/post-sham) affect aperiodic (offset, exponent) and periodic (alpha 8–12 Hz, beta 13–30 Hz) components of the EEG spectra. Specifically, these models included the experimental conditions as fixed effects and participants as random effects to account for inter-individual variability. Pairwise contrasts conducted on post-active vs. post-sham stimulation evaluated their relative differences compared to pre-protocol conditions.

#### 2.4. Results

The LMM analysis of aperiodic components revealed no significant difference in the offset between pre- and post-stimulation (β = 0.02 [-0.05, 0.08], p = 0.614) and between pre- and post-sham (β = 0.01 [-0.06, 0.07], p = 0.888). The relative difference between post-stimulation and post-sham was not significant either (β = 0.02 [-0.05, 0.09], p = 0.629). Similarly, there were no significant changes observed in the exponent between pre- and post-stimulation (β = 0.01 [-0.04, 0.07], p = 0.636), between pre- and post-sham (β = -0.01 [-0.06, 0.05], p = 0.827), and between post-stimulation and post-sham (β = 0.01 [-0.04, 0.06], p = 0.597).

The LMM analysis of periodic components revealed that, in the alpha band (8-12 Hz), the comparison between pre- and post-stimulation was not significant (β = -0.00 [-0.05, 0.04], p = 0.853), but a significant decrease in alpha power was found between pre- and post-sham (β = -0.08 [-0.12, -0.04], p < 0.001). However, the relative difference between post-stimulation and post-sham was itself not significant (β = -0.00 [-0.04, 0.03], p = 0.821). In the beta band (13-30 Hz), the comparison between pre- and post-stimulation was not significant (β = -0.00 [- 0.03, 0.03], p = 0.847), yet a significant decrease in beta power was again observed between pre- and post-sham (β = -0.04 [-0.06, -0.01], p = 0.008). However, the relative difference between post-stimulation and post-sham conditions remained not significant (β = -0.00 [-0.03, 0.03], p = 0.863).

Together, these results suggest that HD-tRNS over the rTPJ during the HDC had no significant influence on either aperiodic or periodic components of post-block eyes-open resting EEG recorded in Experiment 1 (Supplementary Fig. 2), as no measurable differences were observed between pre- and post-stimulation or between post-stimulation and post-sham compared to pre-block recordings.

**Supplementary Figure 2.**
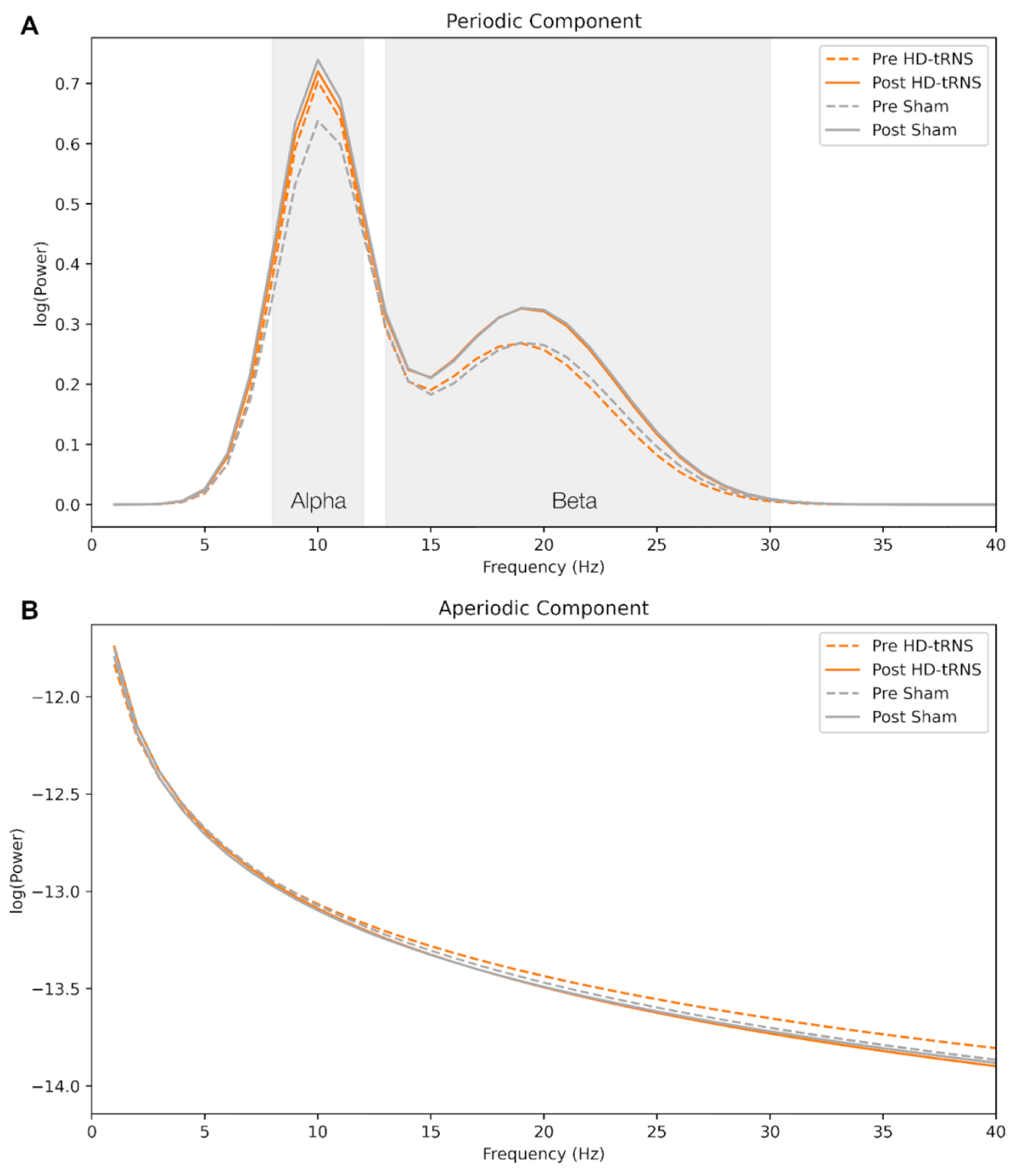
Spectral parametrization of resting-state EEG in (A) periodic and (B) aperiodic components across pre- and post-stimulation/sham in Experiment 1.

### 2.5 Discussion

Despite inducing neuromodulatory effects in social inference, eyes-open resting EEG recorded before and after HDC blocks of Experiment 1 did not reveal any robust changes in periodic (alpha and beta bands) or aperiodic (offset and exponent) components. Crucially, we found no significant differences between post-stimulation and post-sham relative to the pre-task baseline, suggesting that our short, trial-based tRNS sessions did not induce lasting neural changes. This absence of detectable electrophysiological alterations aligns with previous studies indicating that short-duration, online tRNS protocols often produce minimal or no residual effects in resting EEG (Terney et al., 2008). In contrast, research utilizing longer or offline stimulation sessions has reported EEG modulations (Van Doren et al., 2014; Ghin et al., 2021), underscoring that most durable post-stimulation effects on EEG or other neurophysiological markers typically require extended protocols.

Enhanced rTPJ function during active social processing might not necessarily leave enduring resting-state signatures. This is consistent with the idea that tRNS facilitates cognitive processes by interacting with ongoing task-related neural activity rather than inducing specific frequency alterations in EEG (Ke et al., 2024). Consequently, while tRNS can modulate neuronal activity and behavior during task engagement, these effects may not persist once the task concludes, especially under brief stimulation protocols. Furthermore, the low spatial density of our EEG setup (four hybrid electrodes) might have limited our ability to detect subtle or localized neural modulations. High-density EEG recordings, such as those employed by Dumas and colleagues (Dumas et al., 2020) during interaction with the HDC, can provide more detailed insights into ongoing rTPJ activity. However, unlike transcranial alternating current stimulation (tACS), which now benefits from advanced algorithms that can effectively remove linear stimulation artifacts from EEG recordings (Haslacher et al., 2021), tRNS still faces challenges in isolating true neural signals from stimulation-induced random noise during active stimulation. Overall, our protocol likely induced a transient, state-dependent modulation that enhanced rTPJ function during the task but dissipated quickly upon its conclusion. This emphasizes that while short, trial-based tRNS sessions can effectively modulate social inference in real time, these effects do not necessarily translate into lasting changes detectable during resting-state conditions.

### 3. Assessment of HD-tRNS adverse effects

In Experiment 1, four participants experienced moderate fatigue in both blocks, including a single report of moderate heat during the first block. No moderate or severe pain, burn, itch, or metallic taste was reported. In Experiment 2, one participant felt a moderate itch, three reported severe itches (two brief and one lasting for the entire block), two participants reported moderate pain, two experienced moderate or severe burns (which began early and faded by the end), and three reported moderate heat. One participant reported a light metallic taste, and three experienced moderate fatigue. Despite these symptoms and being able to withdraw at any time, all participants chose to complete the experiment.

### 4. Questionnaire of HD-tRNS adverse effects (translated from French)

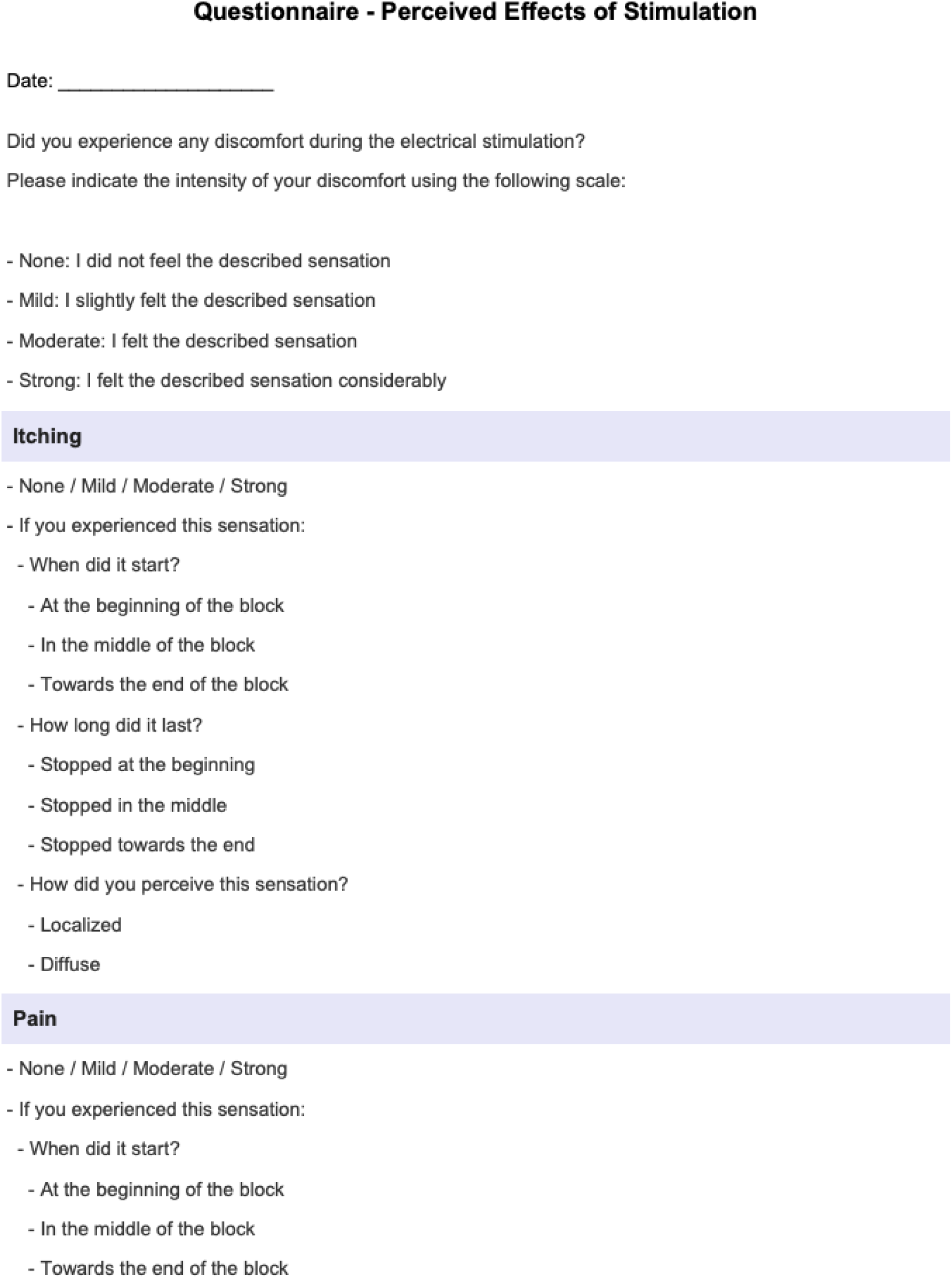
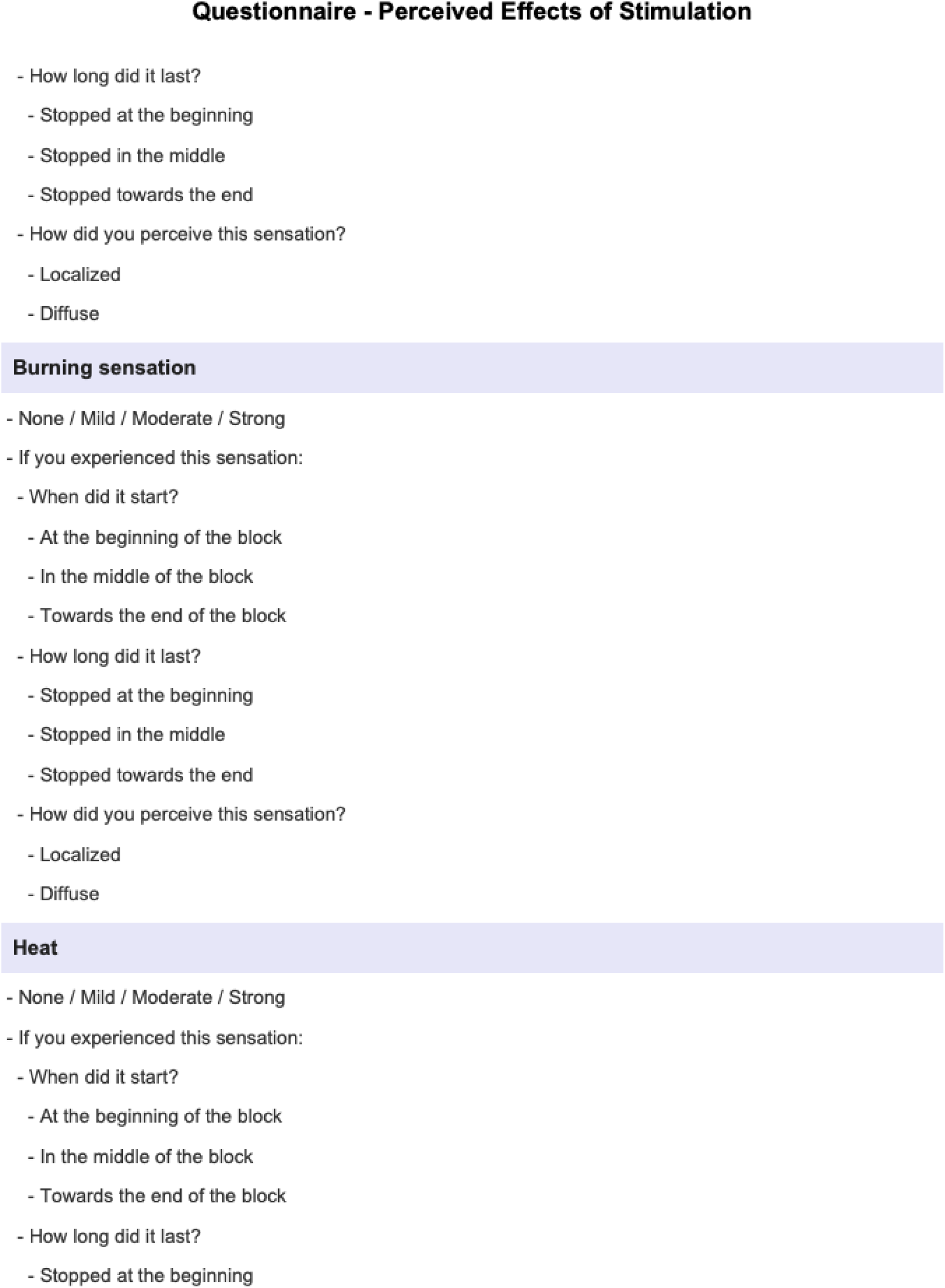
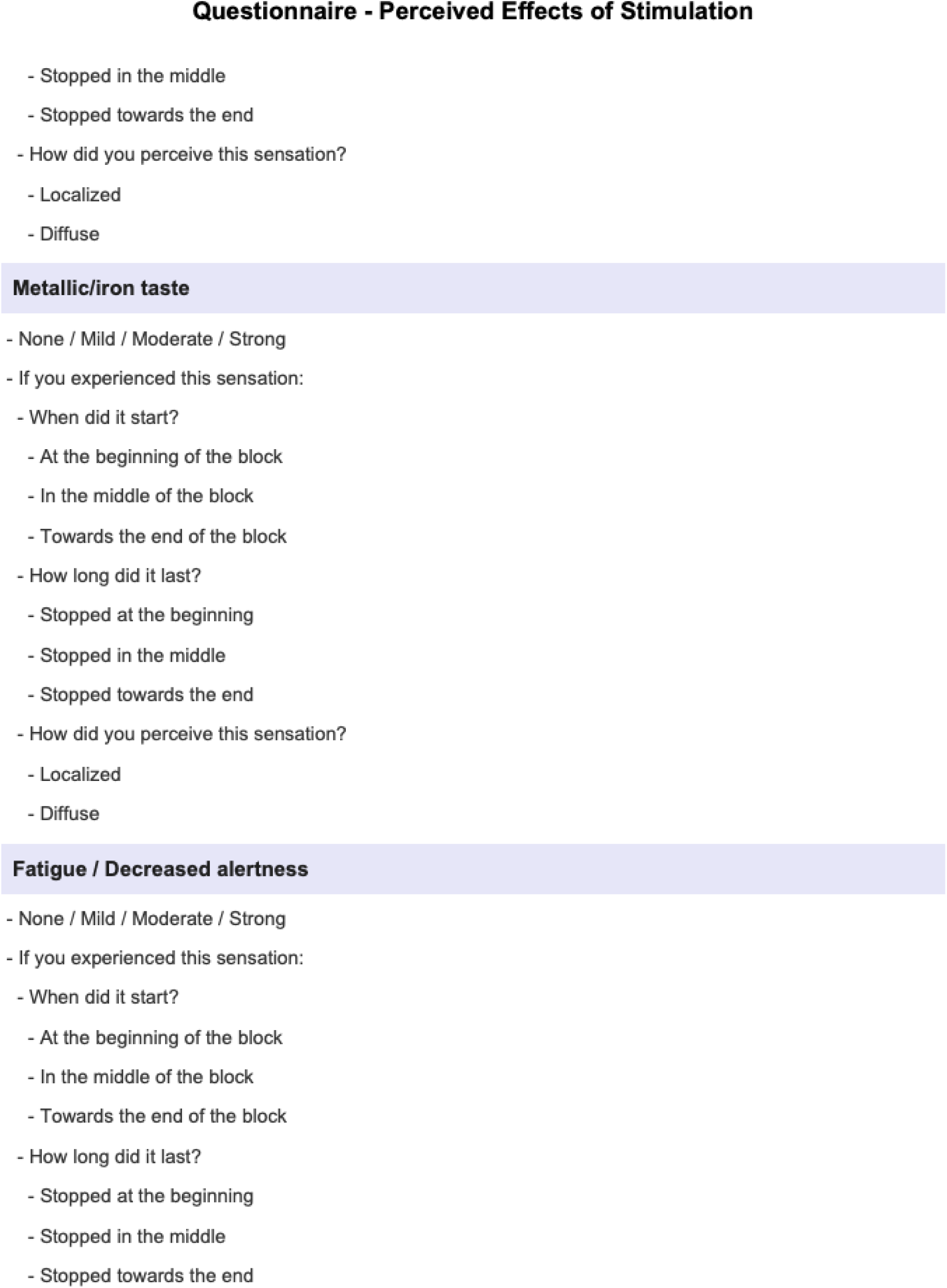
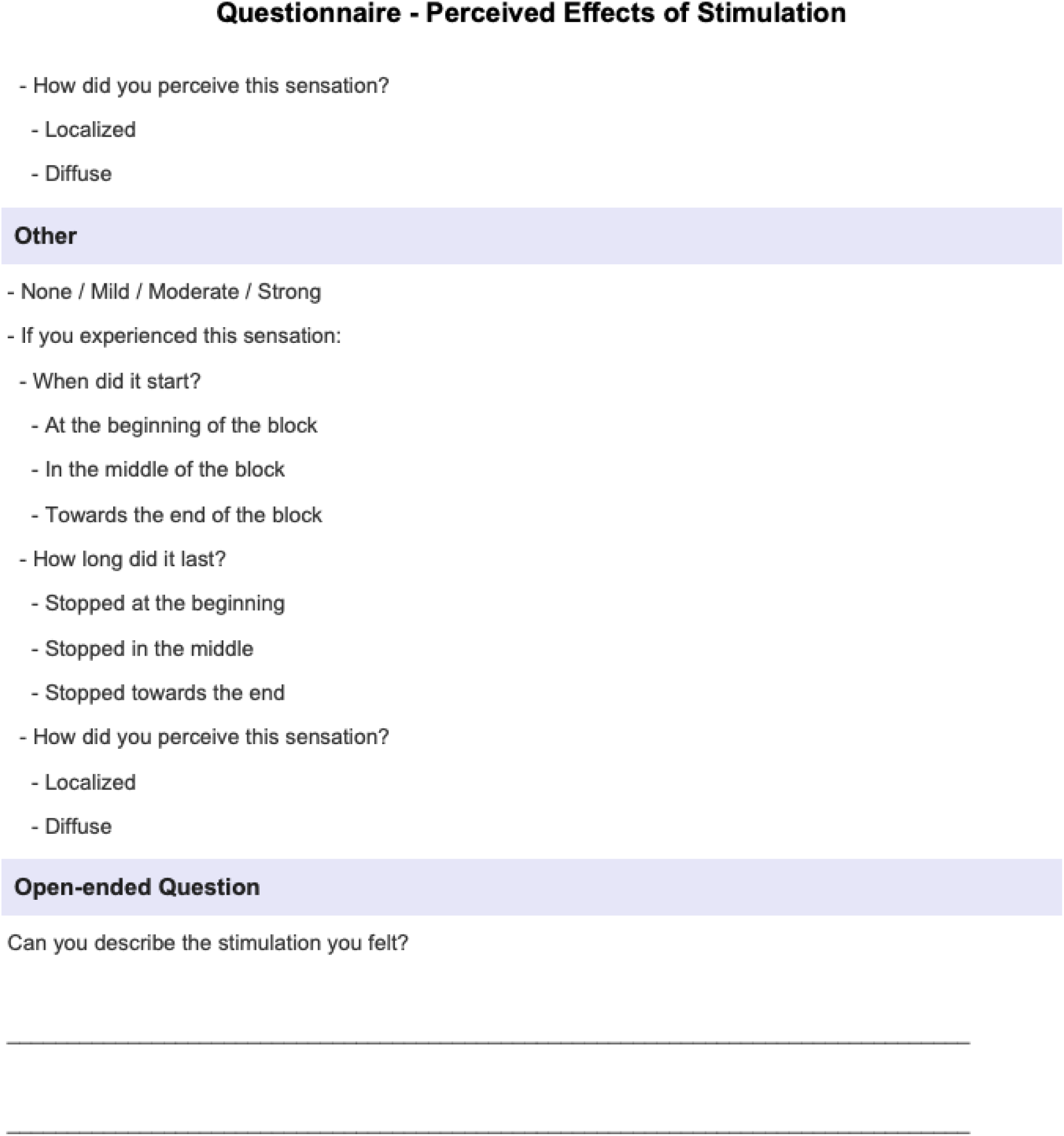

